# A human-specific structural variation at the *ZNF558* locus controls a gene regulatory network during forebrain development

**DOI:** 10.1101/2020.08.18.255562

**Authors:** Pia A. Johansson, Per Ludvik Brattås, Christopher H. Douse, PingHsun Hsieh, Julien Pontis, Daniela Grassi, Raquel Garza, Marie E. Jönsson, Diahann A. M. Atacho, Karolina Pircs, Feride Eren, Yogita Sharma, Jenny Johansson, Didier Trono, Evan E. Eichler, Johan Jakobsson

**Affiliations:** Laboratory of Molecular Neurogenetics, Department of Experimental Medical Science, Wallenberg Neuroscience Center and Lund Stem Cell Center, BMC A11, Lund University, 221 84 Lund, Sweden; Department of Genome Sciences, University of Washington School of Medicine, Seattle, WA, USA 98195; School of Life Sciences, Ecole Polytechnique Fédérale de Lausanne (EPFL), 1015 Lausanne, Switzerland; Howard Hughes Medical Institute, University of Washington, Seattle, WA, USA

**Author notes:** Equal contribution (Pia A. Johansson, Per Ludvik Brattås). Correspondence: Johan Jakobsson, Dept of Experimental Medical Science, Wallenberg Neuroscience Center, BMC A11, 221 84, Lund SWEDEN.

## Abstract

The human forebrain has expanded in size and complexity compared to that of chimpanzee despite limited changes in protein-coding genes, suggesting that gene regulation is an important driver of brain evolution. Here we identify a KRAB-ZFP transcription factor, ZNF558, that is expressed in human but not chimpanzee forebrain neural progenitor cells. ZNF558 evolved as a suppressor of LINE-1 transposons but has been co-opted to regulate the mitophagy gene *SPATA18*, supporting a link between mitochondrial homeostasis and cortical expansion. The unusual on-off switch for *ZNF558* expression resides in a downstream variable number tandem repeat (VNTR) that is contracted in humans relative to chimpanzee. Our data reveal the brain-specific co-option of a transposon-controlling KRAB-ZFP and how a human-specific regulatory network is established by a *cis*-acting structural genome variation. This represents a previously undescribed genetic mechanism in the evolution of the human brain.

## Introduction

The human forebrain has increased in size and complexity after the split between the human and chimpanzee lineages, giving rise to a new level of cognitive functions during hominid evolution (Hill and Walsh, 2005; Lui et al., 2011; Rakic, 2009; Sousa et al., 2017). Cellular and anatomical adaptations have been driven by genetic changes in the human lineage (Enard, 2016), but the actual genetic modifications responsible for this evolutionary process are mostly not understood. Protein-coding genes are highly conserved between human and chimpanzees (Kronenberg et al., 2018). Indeed, aside from the well-studied transcription factor FOXP2 (Lai et al., 2001) there remains limited evidence for a wider impact of amino acid substitutions on human brain evolution. Recently, larger structural variations resulting in gene duplication were implicated in human forebrain function and evolution. *NOTCH2NL*, a human-specific paralog of *NOTCH2*, contributes to cortical development (Fiddes et al., 2018; Suzuki et al., 2018), and duplication of *TBC1D3* and the mitochondrial protein *ARHGAP11B* affected cortical expansion via the basal progenitor populations (Dennis and Eichler, 2016; Florio et al., 2015; Ju et al., 2016; Namba et al., 2020). In addition to amino acid changes and gene innovation, changes in *cis*-regulatory regions have long been thought to contribute to species-specific differences (King and Wilson, 1975), and several studies have revealed divergent gene expression patterns in developing primate brains although their evolutionary impact is unclear (Johnson et al., 2009; Khaitovich et al., 2006; Mora-Bermúdez et al., 2016; Prescott et al., 2015).

One gene family of particular interest in human brain evolution are the Krüppel-associated box (KRAB) domain-containing zinc-finger proteins (KZFPs), the largest individual family of transcription factors in mammalian genomes. KZFPs have undergone a rapid expansion during mammalian and primate evolution and the human genome encodes for at least 350 KZFPs, around 170 of which are primate-specific (Imbeault et al., 2017; Jacobs et al., 2014). Many KZFPs are expressed in the human brain where they are integrated into neuronal gene regulatory networks (Farmiloe et al., 2020; Imbeault et al., 2017). Notably, these expression patterns are different between human and chimpanzee (Nowick et al., 2009). The majority of KZFPs are thought to be transcriptional repressors. Their conserved N-terminal KRAB domain interacts with the epigenetic co-repressor TRIM28, which induces heterochromatin formation and transcriptional repression of targets (Ayyanathan et al., 2003; Emerson and Thomas, 2009; Matsui et al., 2010; Rowe et al., 2010; Sripathy et al., 2006). KZFPs differ mainly in the number and sequence of their zinc finger (ZF) domains, with the number of ZFs ranging between 2 and 40 in humans (Imbeault et al., 2017). The ZFs determine binding specificity: each domain presents a binding surface that is specific for different nucleotide stretches in the target DNA (Patel et al., 2018; Persikov and Singh, 2014). The KZFP family has expanded and diversified through repeated cycles of segmental duplications, giving rise to novel KZFP genes with new targets and biological functions (Nowick et al., 2010).

Specific functions of individual KZFPs have started to be uncovered. Several studies have demonstrated an important role in the repression of transposable elements (TEs) in a variety of cell types, including embryonic stem cells (ESCs) and neural progenitor cells (Brattås et al., 2017; Ecco et al., 2016; Fasching et al., 2015; Najafabadi et al., 2015; Pontis et al., 2019; Rowe and Trono, 2011; Rowe et al., 2010; Turelli et al., 2014; Wolf et al., 2015; Zhang et al., 2019). The rapid expansion of KZFPs in mammalian genomes is correlated with the expansion of TEs, where KZFPs are thought to evolve to target new TE insertions and sequences (Jacobs et al., 2014; Thomas and Schneider, 2011). Indeed, the majority of KZFPs bind to specific families of TEs in human cells (Imbeault et al., 2017; Najafabadi et al., 2015) but it has also been proposed that both KZFPs and TEs have been co-opted by host genomes for broader regulation of transcriptional networks (Brattås et al., 2017; Ecco et al., 2016; Friedli and Trono, 2015; Imbeault et al., 2017).

These observations make KZFPs promising candidates to mediate evolutionary differences between the human and chimpanzee brain. However, there is a lack of experimental data directly addressing this hypothesis. In this study we investigate the co-option of KZFPs in transcriptional regulation during human and chimpanzee brain development. To this end, we established a robust *in vitro* differentiation protocol allowing for quantitative comparisons between forebrain neural progenitor cells (fbNPCs) of human and chimpanzee origin. We discovered several KZFP transcription factors that are highly expressed in human but not chimpanzee fbNPCs. One of these, ZNF558, is a conserved gene that originally evolved to control the expression of LINE-1 elements around 100 million years ago. Our data show that ZNF558 no longer suppresses TEs, but has been co-opted in fbNPCs to regulate a single gene, *SPATA18*, a regulator of mitophagy (Kitamura et al., 2011). Mechanistically, we provide evidence that the unusual on-off switch for ZNF558 expression resides in a downstream variable number tandem repeat (VNTR) that is contracted in humans relative to chimpanzee. Epigenetic manipulation of the human VNTR was sufficient to switch off ZNF558 expression and thereby increase SPATA18 expression. Taken together our data reveal the co-option of a TE-controlling KZFP to regulate a protein-coding gene, and how this regulatory network is controlled by a cis-acting structural genome variation. This finding represents a previously undescribed genetic mechanism for the evolution of the human brain.

## Results

### Derivation of human and chimpanzee forebrain neural progenitor cells

Comparative transcriptomic and epigenetic analyses of chimpanzee and human brain development have been hampered by both availability of material from these species and tissue heterogeneity. However, induced pluripotent stem cells (iPSCs) from chimpanzee and other primates recently became available, enabling *in vitro* modeling studies (Marchetto et al., 2013; Mora-Bermúdez et al., 2016; Romero et al., 2015; Wunderlich et al., 2014). To directly compare human and chimpanzee forebrain neural progenitor cells (fbNPCs), we optimized a defined, feeder-free, 2D differentiation protocol based on dual-SMAD inhibition (Grassi et al., 2020). Chimpanzee and human iPSCs could be maintained *in vitro* under identical conditions (**Fig. 1A**), and upon induction of differentiation we observed a rapid switch to a fbNPC-like morphology in cells from both species (**Fig. 1B**). After two weeks of differentiation we found that both human and chimpanzee fbNPCs expressed high levels of *FOXG1*, a key forebrain marker, while the pluripotency marker *NANOG* was absent from the cultures (**Fig. 1B**, **Supp. Fig. 1A**).

**Figure 1.**
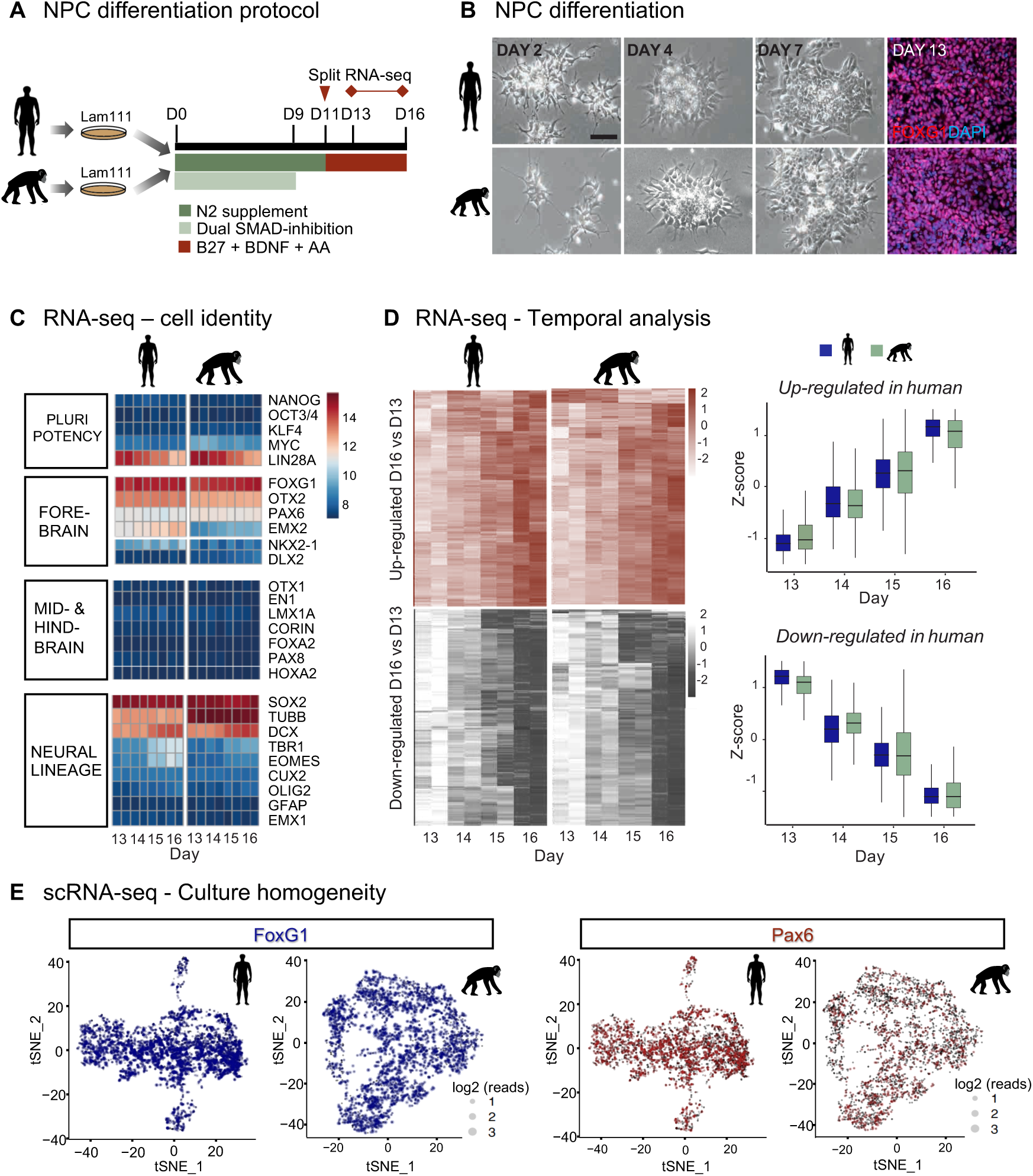
Differentiation of human and chimpanzee iPSCs to forebrain neural progenitor cells. (A) Schematics illustrating the dual-SMAD inhibition-based differentiation protocol from seeding iPSCs at day 0 to harvesting the forebrain progenitors at days 13-16 of differentiation. (B) Brightfield images of human and chimpanzee cells during the first week of differentiation, and FOXG1 immunocytochemistry at day 13, scale bar: 100 μm. (C) Heatmap showing marker expression at days 13-16 of differentiation. (D) Heatmaps (left) displaying genes that are significantly (p adj. < 0.01) up- and down-regulated over time of differentiation (day 16/day 13) in humans, the same set of genes mapped for both species. Boxplots (right) showing the same set of genes. The lower and upper hinges correspond to the first and third quartiles. (E) tSNE-analysis of single-cell RNA-seq data of the forebrain markers PAX6 and FOXG1 in human and chimpanzee fbNPCs.. See also Supp. Fig. S1.

Bulk transcriptome analysis confirmed that both human and chimpanzee fbNPCs expressed appropriate neuronal and forebrain markers, while genes related to other brain regions or other tissues were close to undetectable (**Fig. 1C**). To investigate if human and chimpanzee iPSCs differentiate into fbNPCs with different temporal trajectories e.g. due to differences in cell-cycle progression, we performed bulk RNA-seq at 13, 14, 15 and 16 days of differentiation and analyzed the covariance of gene expression between these time points and the two species. At the selected time points, the fbNPCs corresponded to a differentiation stage just prior to neuronal commitment, demonstrated by the gradual increase in basal progenitor marker *EOMES* and neuronal markers *DCX* and *TBR1* from day 13 to 16 (**Fig. 1C**). We did not see upregulation of glial markers, in line with the stepwise generation of neurons and glia during human brain development (**Fig. 1C**). Globally, a similar set of genes were up- and down-regulated between day 13 and day 16 in human and chimpanzee fbNPCs, indicating that the temporal dynamics of the protocol was closely matched (**Fig. 1D**). We confirmed a limited batch-to-batch variation in the differentiation protocol and the results were consistent in cell lines from different individuals (**Supp. Fig 1B**).

To investigate the heterogeneity of human and chimpanzee fbNPC cultures we performed single-cell RNA-seq analysis at day 14 of differentiation for 4,355 human and 3,620 chimpanzee fbNPCs. Principal component analysis (PCA) showed that 95% of the cells clustered into one major population in both species (**Supp. Fig. 1C-E**). Transcriptional variation within the major population was mainly explained by differences in cell-cycle state rather than cell identity, as these cells clustered into a dense population inseparable on PC1 and PC2 after regressing out cell-cycle effects (**Supp. Fig. 1D**). t-Distributed Stochastic Neighbor Embedding (tSNE) confirmed the presence of a large major population of cells homogeneously expressing the forebrain progenitor markers *FOXG1* and *PAX6* (**Fig. 1E**). The tSNE analysis also revealed two minor populations (<5% of both human and chimpanzee fbNPC cultures), one of which expressed markers associated with early-committed neurons such as *NEUROG1* and *NEUROD1*, while the other related to the endothelial lineage (e.g. *ANKRD1* and *CTGF*) (**Supp. Fig. 1E**). Taken together, immunocytochemistry, bulk and single-cell RNA-seq demonstrated that our 2D differentiation protocol was reproducible and gave rise to temporally- and phenotypically-matched homogeneous cultures of human and chimpanzee fbNPCs, making it a suitable model system for direct comparative analysis.

### Human-specific expression of KZFP transcription factors in forebrain progenitor cells

Next we queried our transcriptomic datasets for differentially expressed genes. We chose to focus on KZFP transcription factors because their evolutionary and biochemical characteristics make them prime candidates for governing species-specific differences in human and chimpanzee brain development. We identified 312 KZFPs that were expressed in at least one sample and 35 that were significantly different between species (27 higher in human, 8 higher in chimp; Wald’s test, *P* < 1.0 × 10^−15^ with Benjamini-Hochberg correction and log_2_-fold-change > 1) (**Fig. 2A**). Seven candidates (*ZNF138, ZNF248, ZNF439, ZNF557, ZNF558, ZNF596* and *ZNF626*) were highly expressed only in human fbNPCs, with nearly no expression in chimpanzee (**Fig. 2B**, **Supp. Fig. 2A**).

**Figure 2.**
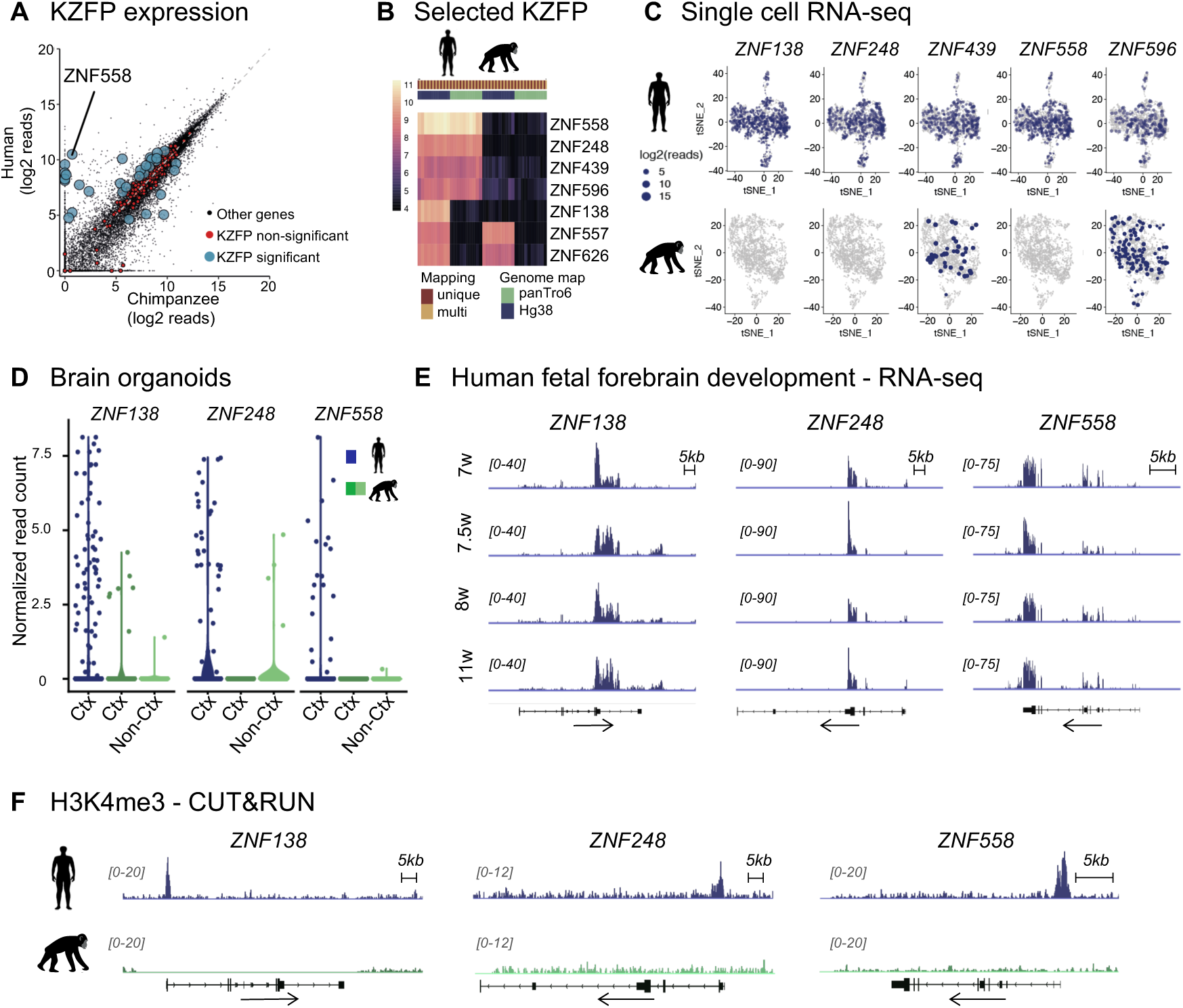
Identification of KZFPs with human-specific expression. (A) Expression levels of genes shown as normalized read counts in human (y-axis) and chimp (x-axis) in fbNPCs. KZFP genes are marked red if not significantly different and marked blue if significantly different (P < 1e-15; Benjamini-Hochberg corrected, and log2-fold change > 1) between human and chimp. (B) Heatmap of seven KZFPs with high expression in human and no expression in chimp and mapping of all samples to both the human and chimp reference genome, both with unique and multi-mapping approaches. (C) tSNE plots with visualization of expression of KZFPs in single cell sequencing data of humanand chimp fbNPCs. Dots represent cells with no reads mapped to the gene, size of dots proportional to level of expression of gene. (D) Normalized read counts of KZFP expression in single cell sequencing data of human (blue) and chimp (green) cortical organoid data(Mora-Bermúdez et al., 2016). Each dot represents one cell. Ctx: Cortical-like cells. Non-Ctx: Non cortical cells, based on marker expression and clustering. (E) RNA-seq analysis of KZFP expression in dissected forebrain tissue from aborted human embryos of different gestational age (w, weeks). Scale is RPKM (reads per kilobase per million mapped reads). (F) Normalized CUT&RUN coverage tracks illustrating H3K4me3 levels over KZFP genes in human (top, blue) and chimp (bottom, green) fbNPCs. Data were obtained in biological duplicate (i.e. cells derived from two individuals for each species) with similar results. CUT&RUN data were normalized based on spike-in DNA (see methods). See also Supp. Fig 2.

The massive expansion of the KZFP family in mammals has arisen through rounds of segmental duplication events, making them a challenging gene family to study with comparative genomic analyses. To confirm that human-specific expression was not caused by biases in mapping, reference genome builds or gene annotation, we mapped all samples to both human (GRCh38) and chimpanzee (PanTro6) assemblies. *ZNF138* expression was only detected when mapping human samples to GRCh38, but not when mapping human samples to PanTro6, nor in chimp samples mapping to either assembly. Pairwise alignment of human and chimp *ZNF138* coding sequences revealed only 34% sequence identity, with point mutations and several deletions in the human sequence – explaining why reads from human samples do not map to the chimpanzee genome. We conclude that *ZNF138* has diverged in both sequence and expression pattern in forebrain progenitors. *ZNF557* and *ZNF626* were detected in chimp samples only when mapping to GRCh38. This suggested that issues with the PanTro6 assembly prevented mapping of these transcripts, invalidating attempts to infer human-specific expression of these genes. Four candidate genes (*ZNF248*, *ZNF439*, *ZNF558* and *ZNF596*) were exclusively expressed in human samples, despite high mappability for both human and chimp orthologues (**Fig. 2B**).

To further dissect the expression divergence for the five candidate KZFPs (*ZNF138*, *ZNF248*, *ZNF439*, *ZNF558* and *ZNF596*) with human-specific expression in the developing forebrain, we used our single-cell RNA-seq analysis of human and chimp fbNPCs. This analysis confirmed that all five candidates were highly expressed in human samples (**Fig. 2C**). *ZNF138*, *ZNF248* and *ZNF558* were exclusively expressed in human cells, while *ZNF439* and *ZNF596* were also detected in a rare number of chimp cells (**Fig. 2C**, **Supp. Fig. 2B**). The candidates were expressed throughout the cell population, excluding the possibility that the human-specific expression was due to a specific sub-population in the human samples.

To confirm that human-specific expression of candidate KZFPs was not an *in vitro* artefact originating from our cell culture system or the low number of individuals, we analyzed expression patterns in human and chimp cerebral organoids from different labs using iPSCs derived from different individuals. This analysis confirmed that *ZNF138*, *ZNF248* and *ZNF558* are exclusively expressed in human cells (Field et al., 2019; Kanton et al., 2019; Mora-Bermúdez et al., 2016) (**Fig. 2D, Supp Fig. 2C,D**), and RNA-seq data from human fetal forebrain samples confirmed their expression during human forebrain development (**Fig. 2E**). Finally, CUT&RUN epigenomic profiling in human and chimp fbNPCs demonstrated a striking human-specific enrichment of the activating epigenetic mark H3K4me3 over the promoters of the candidate genes, confirming that the observed differences in RNA levels results from differences in transcriptional activity (**Fig. 2F**). Taken together, our analysis identified three KZFPs expressed in human but not chimp fbNPCs (*ZNF138*, *ZNF248* and *ZNF558*). We selected *ZNF558* for further analysis as it was exclusively and highly expressed in human cells in all samples investigated.

### *ZNF558* has evolved under constraint in the mammalian lineage

Many KZFPs have undergone rapid evolution in mammalian and primate evolution, where new genes arise and diversify through segmental duplications (Nowick et al., 2010). However, *ZNF558* is estimated to have originated before the common ancestor of afrotherian mammals, about 105 million years ago (Mya), with orthologues in a range of placental mammals (**Fig. 3A**). A duplication event occurred in the last common ancestor with new world monkeys, giving rise to a paralogue (*ZNF557*), which was expressed in both human and chimpanzee fbNPC samples. *ZNF557* encodes one more ZF domain (10 in total) than *ZNF558*, and 8 of the 9 common ZF domains have one or more non-synonymous mutations in DNA binding residues relative to ZNF558, suggesting that positive selection acted to diversify function of these paralogues resulting in divergent DNA targets.

**Figure 3.**
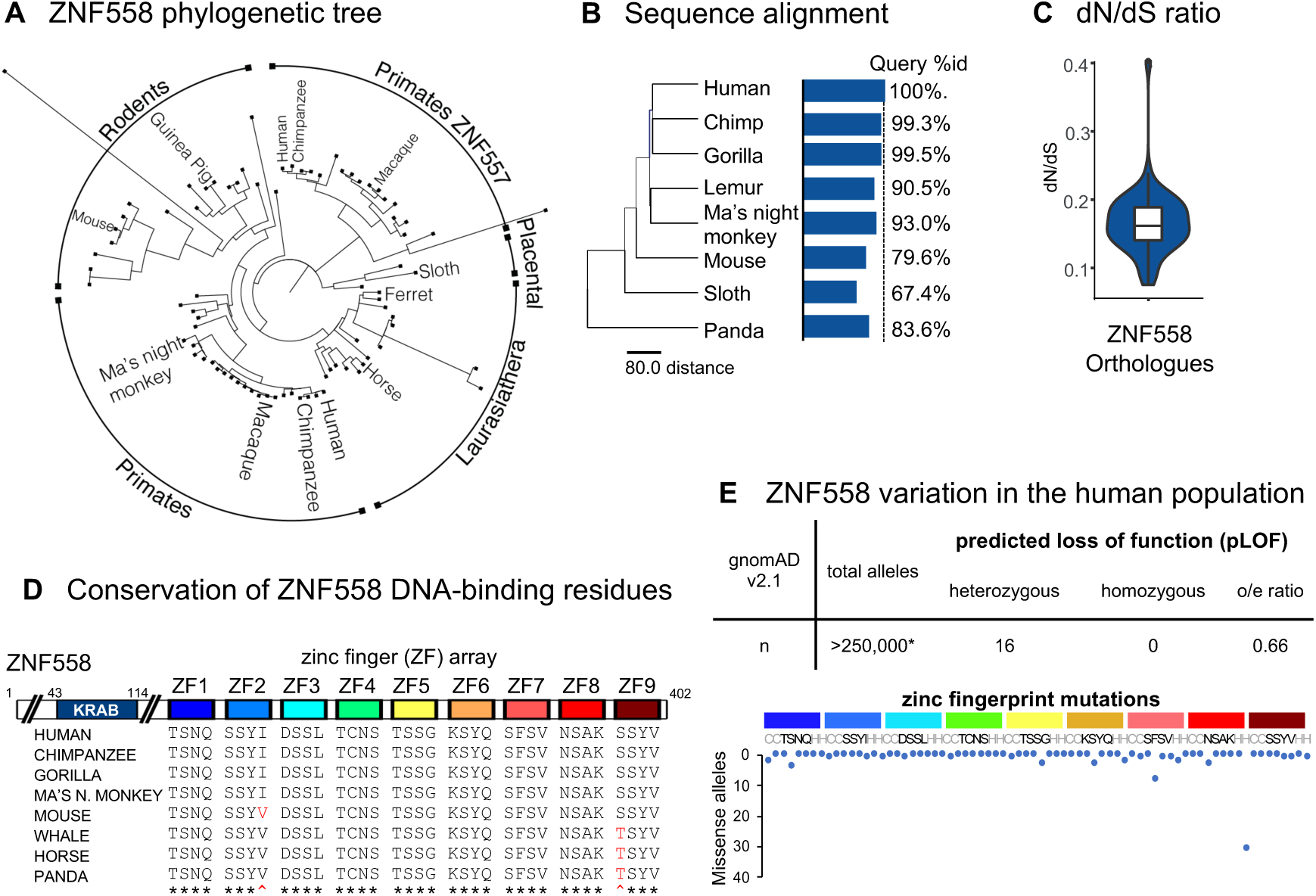
Evolutionary history of ZNF558. (A) Phylogenetic tree of ZNF558 orthologues and paralogues (ZNF557). (B) Percentage of orthologous protein sequence matching human ZNF558 protein sequence in selected species. (C) dN/dS ratio of all ZNF558 orthologues identified in Ensembl. (D) Domain structure of ZNF558 (top). Multi-sequence alignment of DNA-contacting residues in the zinc finger (ZF) array of selected ZNF558 orthologues (bottom). The four amino acids shown per ZF are defined as the −1,−4,−5,−7 positions relative to the first histidine of each C2H2 zinc finger. (E) Genetic variation of *ZNF558* in the human population. Shown are the number of high-quality genotypes annotated as predicted loss of function (pLOF, top) or leading to missense substitutions in zinc- (grey) or DNA-binding (black) resides from ZF domains (bottom). o/e, observed to expected ratio of pLOF variants. Data from gnomAD v2.1 (Karczewski et al., 2020).

Pairwise alignments revealed high conservation among detected ZNF558 orthologues (**Fig. 3B**). Analysis of the ratio between non-synonymous to synonymous mutations (dN/dS) suggests that ZNF558 has evolved under constraint in the mammalian lineage, with a median dN/dS ratio of 0.159 (**Fig. 3C**). KZFP DNA binding is mediated by tandem C2H2 zinc finger (ZF) domains, each of which interacts with contiguous stretches of 2-4 nucleotides per finger (Patel et al., 2018; Persikov and Singh, 2014). In particular, the identities of four residues in each ZF are responsible for sequence specificity, with multiple ZF domains combining to encode a particular ‘zinc fingerprint’ (Imbeault et al., 2017). To test whether the DNA-contacting residues of its 9 ZFs have evolved during recent mammalian evolution, we aligned sequences of the ZNF558 fingerprint among eight mammals and found that these residues are almost completely conserved. All primates display identical binding residues, while single substitutions in mouse, whale and panda proteins are conservative (Ile to Val; Ser to Thr) and thus unlikely to have a major effect on DNA binding (**Fig. 3D**).

We next investigated the presence of *ZNF558* gene variants in the human population using a database (gnomAD v2) of 125,748 exome and 15,708 whole-genome sequences from unrelated individuals (Karczewski et al., 2020). Heterozygous predicted loss of function (pLoF) alleles were found at slightly below the expected frequency (depth-corrected observed/expected ratio oe = 0.66; 90% CI 0.48-1.12), and no individuals harboring homozygous pLoF variants were identified (**Fig. 3E**). Analysis of missense variants showed that the KRAB domain is strictly conserved in putative TRIM28-interacting residues (Friedman et al., 1996; Murphy et al., 2016). We also found low variant counts in the zinc fingerprints and zinc-coordinating residues among the human population, with one exception, a H370P substitution in ZF8 found in about 1 in 1000 Finnish individuals (**Fig. 3E**). Taken together, this evolutionary and population-wide analysis of ZNF558 conservation and variation indicate that the protein has been under stringent evolutionary constraint for around 100 million years, in line with an important role in mammalian and human physiology.

### ZNF558 binds evolutionarily old LINE-1 elements and protein-coding genes

The expansion of KZFPs in mammalian genomes is thought to be driven by adaptation to rapidly-evolving TEs (Jacobs et al., 2014; Thomas and Schneider, 2011). It was therefore striking that our candidate gene for human/chimpanzee divergence, *ZNF558*, is highly conserved. This suggests that it may have been co-opted for regulating non-TE targets early in mammalian evolution. To investigate the function of ZNF558, we first considered the DNA binding preferences of its nine-membered zinc finger (ZF) array. Chromatin immunoprecipitation with exonuclease digestion (ChIP–exo) data on HA-tagged ZNF558 expressing 293T cells showed robust signal enrichment and, notably, no overlap in target specificity between ZNF558 and its paralogue ZNF557 (Imbeault et al., 2017)(**Fig. 4A**). Computational modeling of the ZNF558 binding motif with overlapping 4-nt subsites suggested a T-rich binding sequence of 28 nucleotides. We found that the experimental binding motif partly matched the prediction with some gaps (**Fig. 4B**). Longer arrays are known to deviate more from the canonical binding model and A/T-rich sites are less predictable, so such discrepancies were not unexpected (Patel et al., 2018).

**Figure 4.**
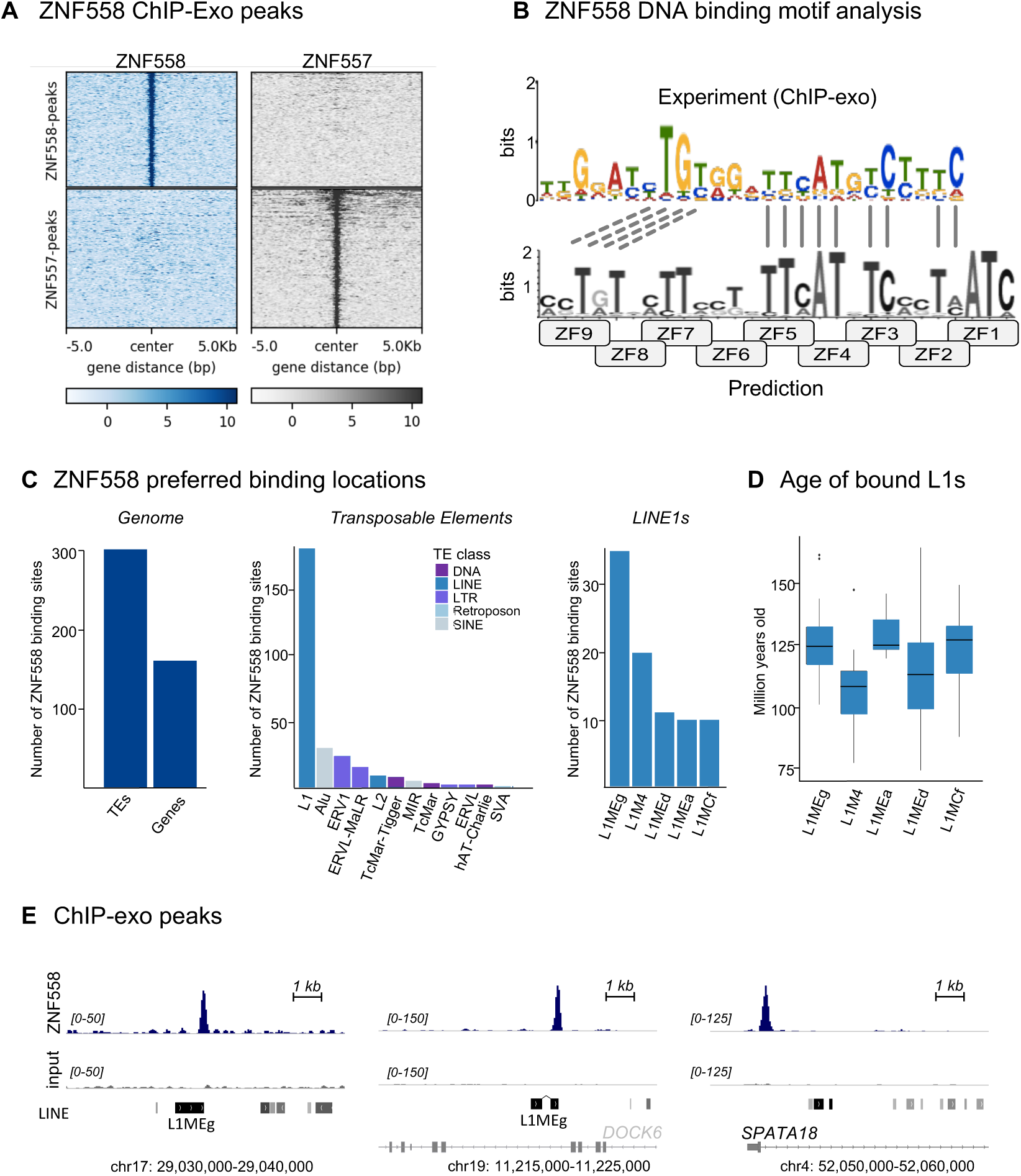
ZNF558 genome binding analysis. (A) Heatmaps of ZNF558 and ZNF557 ChIP signal [REF pmid 28273063] plotted over high-confidence ZNF558 (top) and ZNF557 (bottom) peaks. (B) ZNF558 binding motif analysis based on computational prediction and experimental ChIP-exo data. (C) ZNF558 binding sites in the human genome (left), TEs (middle) and L1-subfamilies (right). Note that TE overlaps include both intergenic and intronic TEs. (D) Evolutionary age of the top five ZNF558-bound L1 families. (E) Example 10-kb screenshots of ZNF558 ChIP-exo peaks. Shown are instances of intergenic and intronic L1MEg (left, middle) and gene targets (right). Scales are RPKM, reads per kilobase per million mapped reads.

We next investigated the features of genomic regions bound by ZNF558. We found that 54% of the 465 high-confidence sites were located in TEs, of which the majority were LINE-1 elements (**Fig. 4C**). Out of these ZNF558-bound L1 elements, the majority of the binding sites were located in evolutionary-old L1 families such as L1MEg, L1M4 and L1Med (**Fig. 4C-E**). These L1 families are remnants of old transposition events and have since degenerated and lost transposition capacity. Estimation of the evolutionary age of the ZNF558-bound elements revealed that the majority were dated around 100 million years old, which correlates well with the age of *ZNF558* (**Fig. 4D**). This observation is in line with a model where ZNF558 originally evolved to repress then-active L1 elements. These L1s would then have degenerated and ZNF558 could have been co-opted to control other non-TE genomic targets. In agreement with this model, we noted that several of the ZNF558 target sites overlap with protein-coding genes (**Fig. 4C**). For example, the fourth highest-scoring ChIP-exo peak was located just downstream of the first exon of *SPATA18*, a gene involved in mitochondrial homeostasis (**Fig. 4E**). These co-opted ZNF558 targets may be important for mammalian physiology, since ZNF558 has been under stringent positive selection for a long time. If this scenario is true, then the L1 binding that remains today likely represents a genomic fossil.

### CRISPRi-mediated transcriptional silencing reveals *SPATA18* as the functional target of ZNF558

To investigate the functional relevance of the ZNF558 binding sites in human fbNPCs, we designed a CRISPR inhibition (CRISPRi) strategy to silence ZNF558 expression (**Fig. 5A**). We targeted two distinct guide RNAs (gRNA) to a genomic region located next to the *ZNF558* transcription start site and co-expressed these with a transcriptional repressor domain fused to catalytically-dead Cas9 (dCas9). Transduction of human iPSCs resulted in efficient silencing of *ZNF558* upon differentiation to fbNPCs (**Fig. 5A-B**), but no difference in differentiation capacity or expression of cell fate markers compared to controls (**Supp. Fig. 3A-B**).

**Figure 5.**
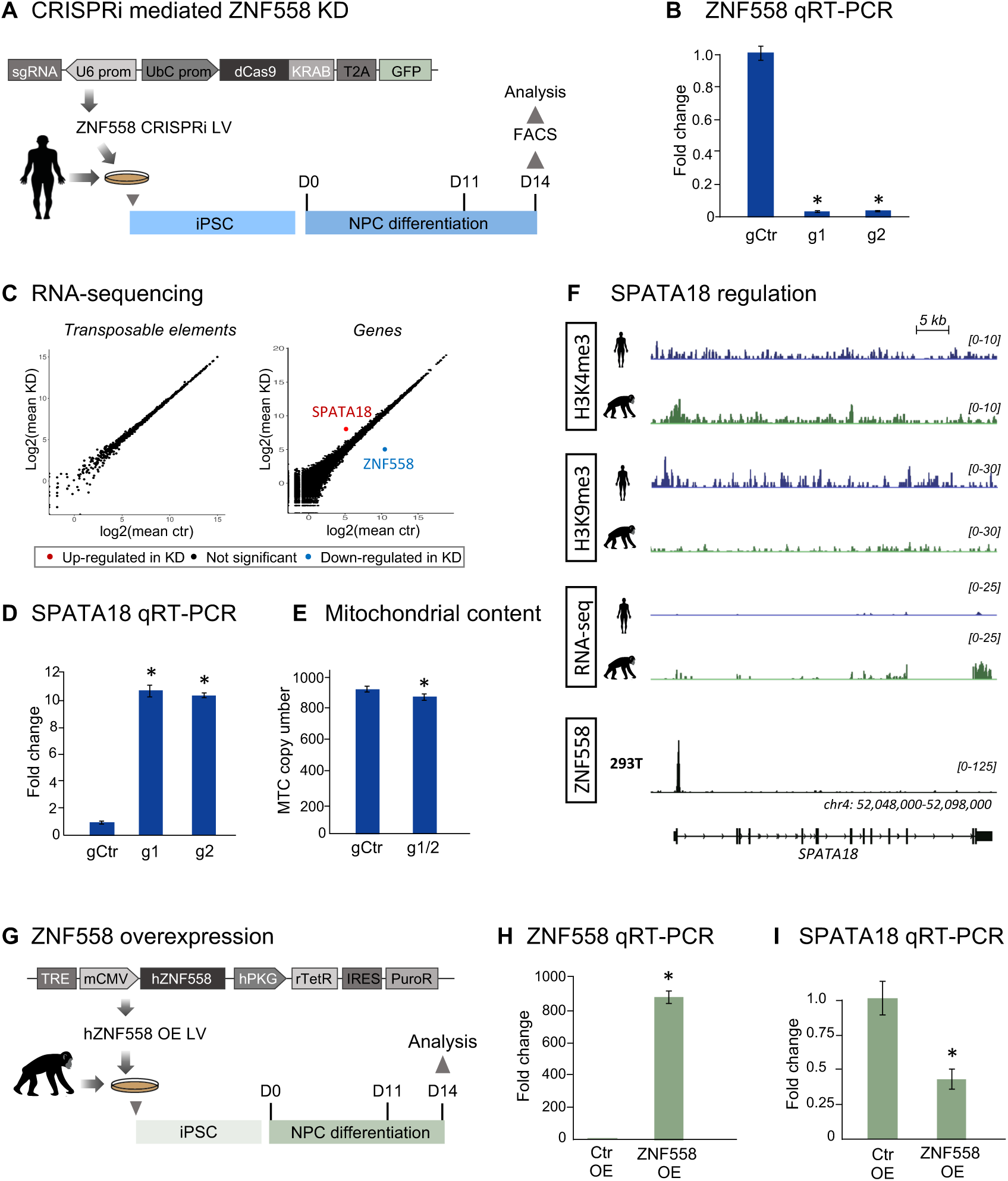
Functional analysis of ZNF558 in fbNPCs. (A) Schematics illustrating the CRISPRi-based strategy to silence ZNF558 expression in human fbNPCs. iPSCs were transduced with a lentiviral CRISPRi construct and then differentiated for 14 days before analysis. (B) qRT-PCR analysis of ZNF558 expression in fbNPCs after CRISPRi-silencing (n=4), p-adj <0.05, log2(fc)>0. (C) RNA-seq analysis of ZNF558-silenced fbNPCs for expression of TEs (left) and protein coding genes (right). (D) qRT-PCR analysis of SPATA18 expression in fbNPCs after CRISPRi-silencing (n=4). (E) PCR of mitochondrial DNA in fbNPCs after ZNF558 CRISPRi-silencing (n=4-6). (F) Normalized epigenomic (H3K4me3, H3K9me3), transcriptomic (RNA-seq) and ZNF558 binding (ChIP-exo) analysis of the *SPATA18* locus in human and chimp fbNPCs. Apart from the ChIP-exo analysis in HEK293T (Imbeault et al., 2017), data were obtained in at least biological duplicate (i.e. cells derived from two individuals for each species) with similar results. CUT&RUN data were normalized based on spike-in DNA (see methods). Scales for RNA-seq and ChIP-exo data are RPKM, reads per kilobase per million mapped reads. (G) Schematics illustrating the experimental strategy to overexpress ZNF558 in chimpanzee fbNPCs. (H-I) qRT-PCR analysis of ZNF558 and SPATA18 in chimpanzee fbNPCs transduced with a lentiviral vector expressing ZNF558 (n=4). * = p<0.05 (Student’s T-test), error bars are mean +/− s.e.m. See also Supp. Fig. 3.

Next we performed RNA-seq on *ZNF558*-silenced fbNPCs and analyzed the transcriptome for alterations in TE and gene expression. Remarkably we found a single protein-coding gene – *SPATA18 –* to be upregulated upon ZNF558 silencing (**Fig. 5C**-**D**, **Supp. Fig. 3C, F**), but no alterations in TE expression either when querying expression of unique elements or entire families. These data demonstrate that ZNF558 has been co-opted in human fbNPCs to regulate *SPATA18*, and that its binding sites in old LINE-1 elements do not have detectable consequences on transcription of those TEs.

### ZNF558 regulates SPATA18 expression in human and chimp fbNPCs

Next we considered the species-specificity of *SPATA18* repression by ZNF558. Since transcription factors and their DNA target sites co-evolve, we analyzed the conservation of the ZNF558 binding site in a range of mammalian *SPATA18* genes and found that it is conserved only in primates (**Supp. Fig. 3E**). The conservation of the binding site in chimpanzee *SPATA18* suggests that the lack of ZNF558 expression in chimpanzee fbNPCs should result in increased expression of SPATA18 in these cells. In accordance with this model, epigenomic profiles over the *SPATA18* promoter in chimpanzee fbNPCs revealed enrichment of H3K4me3 but not H3K9me3 with concomitant expression of SPATA18 RNA (**Fig. 5F**). RNA-seq analysis showed a 2.4-fold increase in chimpanzee compared to human fNPCs. Increased *SPATA18* expression in chimpanzee was also observed in cerebral organoids (Field et al., 2019)(**Supp. Fig. 3D**). To further test that ZNF558 has the capacity to regulate *SPATA18* in chimpanzee fbNPCS, we transduced chimp iPSCs with a lentiviral human ZNF558 overexpression construct (**Fig. 5G-H**). Following differentiation to fbNPCs, *SPATA18* was significantly downregulated upon ZNF558 overexpression (**Fig. 5H-I**, **Supp. Fig. 3G**).

Together, our results demonstrate that the functional consequence of differential *ZNF558* expression in human and chimp forebrain development is the repression of a single gene, *SPATA18*. The product of *SPATA18* is mitochondrial eating protein (MIEAP), a mitophagy regulator (Kitamura et al., 2011). CRISPRi-mediated ZNF558 silencing resulted in a small but significant decrease in mitochondrial content in human fbNPCs (**Fig. 5E**) in line with upregulated SPATA18 levels perturbing mitochondrial homeostasis due to excessive mitophagy. It is noteworthy that differences in mitochondrial homeostasis have recently been associated with human-specific cortical expansion through regulation of forebrain basal progenitors (Namba et al., 2020).

### Regulation of *ZNF558* is controlled by a *cis*-acting VNTR

The difference in *SPATA18* expression between human and chimpanzee fbNPCs is a result of upstream changes in *ZNF558* expression. Given that *ZNF558* is conserved between human and chimpanzee, we next explored the underlying genetic basis for differential expression between the two species. We found limited variation in the human and chimpanzee genomic context upstream of *ZNF558*, suggesting that alterations of promoter structure are unlikely to be responsible. However, downstream of *ZNF558* we noted a large repetitive region in the human genome assembly (chr19:8735220-8794192, GRCh38, **Fig. 6A**). We investigated copy number variation at this locus by generating read-depth profiles for a collection of 1,112 high-coverage human and non-human great ape genomes from publicly available datasets (**Fig. 6B**). Our analysis identified a variable tandem repeat (VNTR) motif with unit length 7,460 bp, which showed striking copy number variation across populations and species (**Fig. 6B,C**). In particular, we observed a significant difference in copy number at this VNTR locus between the human and non-human primate samples: all non-human great ape samples except for orangutans, have significantly higher copy numbers (>70) of this VNTR compared to humans (mean: 24-43 copies) (*P* = 8.61 × 10^−21^, Mann-Whitney *U* test, **Fig. 6C**). This result is consistent with qPCR data using genomic DNA obtained from the human and chimpanzee iPSC lines used in the current study (**Fig. 6D**). One interesting outlier in this analysis is the orangutan, which has a VNTR copy number similar to that of humans (mean 36, standard deviation 8.1) compared to that of other non-human great apes. Published transcriptome data showed high ZNF558 transcript levels in orangutan forebrain organoids (Field et al., 2019)(**Supp. Fig. 2D**). Together these results suggest that lower VNTR copy number correlates with higher ZNF558 expression. We therefore hypothesized that this repetitive genomic region downstream of *ZNF558* is responsible for its differential expression.

**Figure 6.**
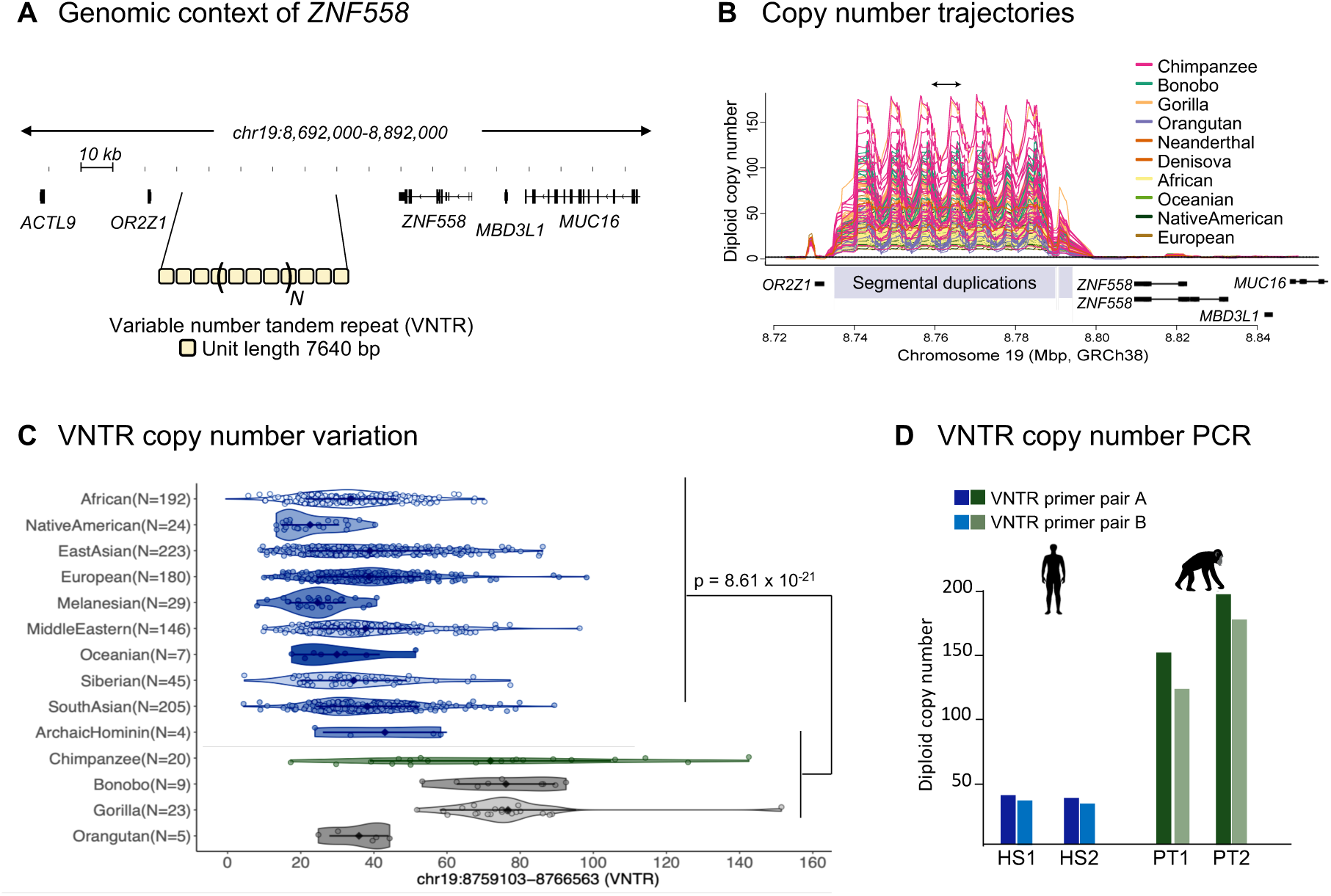
Copy number differences at a VNTR locus downstream of *ZNF558* among humans and great apes. (A) Schematic illustrating the presence of a VNTR downstream of the *ZNF558* gene on chromosome 19 in the GRCh38 assembly. (B) Copy number trajectories illustrating an increase in copy number for a 7.46-kbp VNTR motif (chr19:8,759,103−8,766,563, GRCh38) in chimpanzee, bonobo, and gorilla relative to humans. Each line is the inferred copy number trajectory of a given sample over this region. Black arrows at the top indicate a unit of the VNTR motif. The dashed lines indicate diploid copy number two. (C) Copy number variation for the 7.46-kbp VNTR motif among archaic and modern humans and non-human great ape samples. Each dot is the overall (diploid) copy number for a given sample. Black diamonds and bars represent the mean and one-standard deviation of the VNTR copy numbers in individual populations and species. The p value for copy number differentiation is computed using the Mann-Whitney *U* test. (E) qPCR analysis of the VNTR copy number in the human and chimp iPSC-lines used in this study.

### Epigenetic manipulation of the human VNTR switches the ZNF558-SPATA18 regulatory network

Finally, we investigated the mechanism of *cis-*regulation by the VNTR. We considered that the longer VNTR in chimpanzee could have established a repressive hub leading to *ZNF558* silencing. Profiling of H3K27me3 and H3K9me3 levels over the human and chimp VNTRs is challenging due to the low mappability of this genomic region. However, exploiting spike-in normalization of CUT&RUN data (see methods), we confirmed that relative read depths for control IgG tracks matched relative VNTR copy numbers (**Supp. Fig. 3H**), thereby validating our mapping strategy. H3K9me3 but not H3K27me3 enrichment was detected over the VNTR in both human and chimp samples using this approach (**Fig 7A,B**). These data are consistent with a model in which the VNTR attracts transcriptionally-repressive chromatin mark H3K9me3, the functional consequence of which is controlled by the size of the VNTR. Indeed, RNA-seq analysis showed evidence of transcription within the VNTR in human but not chimp fbNPCs (**Fig. 7A**). Notably, we also found evidence of a human-specific long non-coding RNA originating in the VNTR transcribed in the antisense orientation relative to *ZNF558*, which could be confirmed with qRT-PCR analysis (**Supp. Fig 3I**). These observations suggest that the shorter human VNTR is not fully silenced.

**Figure 7.**
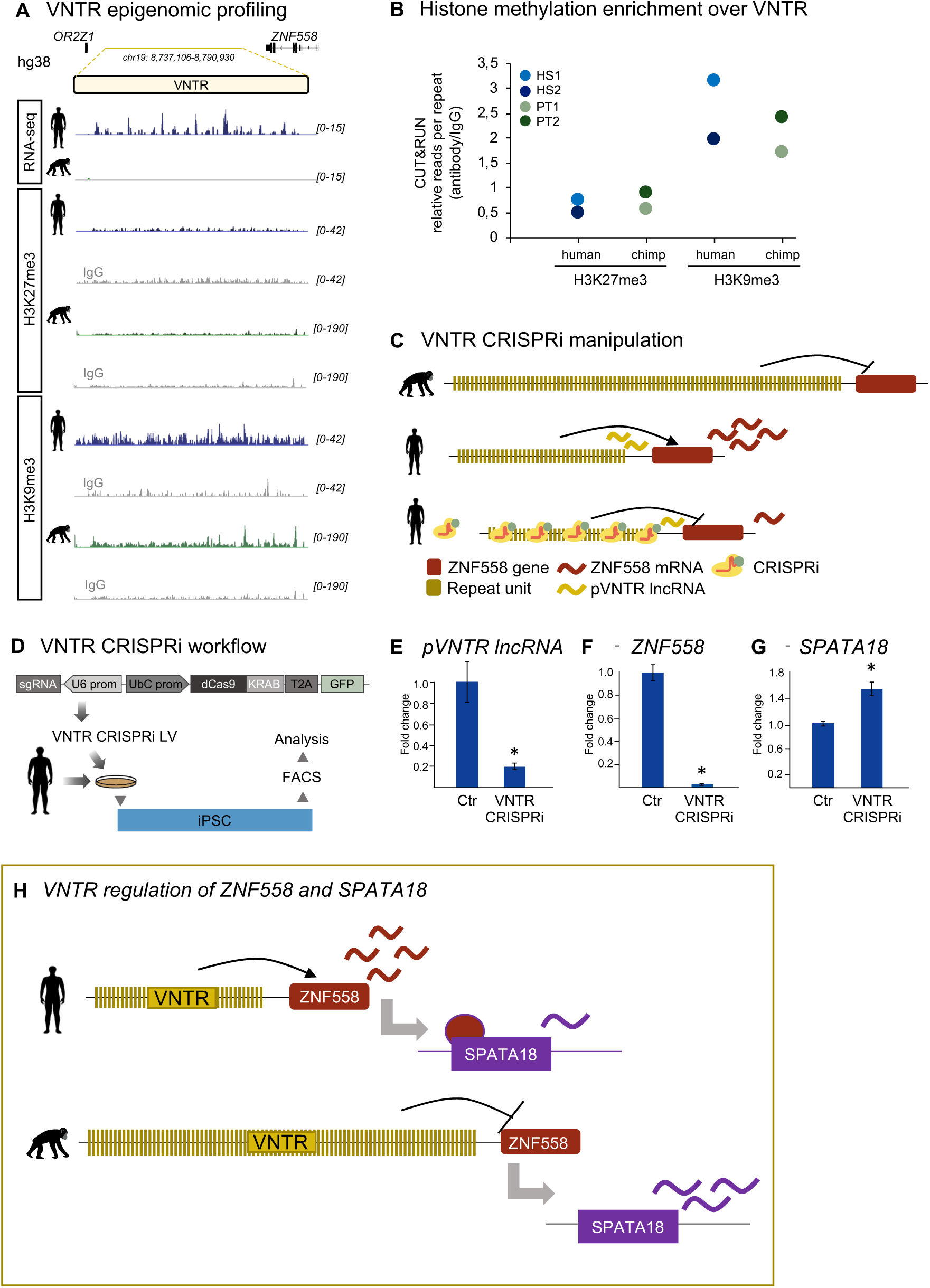
A downstream VNTR controls ZNF558 expression in *cis*. (A) Normalized epigenomic (H3K9me3, H3K27me3) and transcriptomic (RNA-seq) analysis of the VNTR downstream of *ZNF558* in human and chimp fbNPCs. Data were obtained in at least biological duplicate (i.e. cells derived from two individuals for each species) with similar results. CUT&RUN data were normalized based on spike-in DNA (see methods). Scales for RNA-seq are RPKM, reads per kilobase per million mapped reads. (B) Relative CUT&RUN read counts (antibody/IgG control) over the VNTR in human and chimp fbNPCs for repressive chromatin marks H3K27me3 and H3K9me3. Data shown are two biological replicates. IgG controls were performed for each experiment. (C-D) Schematic illustrating the CRISPRi-based multi-guide strategy to epigenetically silence the human VNTR. pVNTR lncRNA = human-specific putative lncRNA originating from within in the VNTR extending towards ZNF558. (E-G) qRT-PCR analysis of VNTR-lncRNA, ZNF558 and SPATA18 in human iPSCs (HS1 and HS2) transduced with VNTR-CRISPRi lentiviral vectors (n=5-6). * = p<0.05, (Student’s t-test), data represented as mean +/− s.e.m. (H) Illustration of the key findings in this study visualizing how the shortening of the VNTR in humans results in loss of epigenetic silencing. The downstream effect is transcriptional activation of ZNF558 activation resulting in SPATA18 repression. See also Supp. Fig. 3.

To experimentally test the model of VNTR-mediated *cis-*repression, we employed a multi-guide CRISPRi strategy to impose silencing over the shorter human VNTR. We designed three different gRNAs to target different parts of the VNTR unit to co-express with the dCas9 repressor fusion (**Fig. 7C,D**). Upon iPSC transduction, we observed reduced expression of the VNTR-associated lncRNA (**Fig. 7E**), in line with targeted silencing. Strikingly, we observed robust inhibition of ZNF558 transcription upon VNTR silencing, which in turn led to increased levels of SPATA18 transcripts (**Fig. 7F-G**). Together these data support the model of a brain-specific gene-regulatory network in which a downstream VNTR controls *ZNF558* expression in *cis*, which in turn affects repression of its target gene *SPATA18* (**Fig. 7H**).

## Discussion

In this study we show how genetic alterations in non-coding regions of the genome can control the activity of conserved protein-coding genes, resulting in the establishment of species-specific transcriptional networks. To date there is limited evidence that changes in *cis*-acting regulatory elements are important for human brain evolution. Classically, non-coding genetic changes are thought to result in gene expression differences along a gradient, leading to slightly more or less mRNA-product of a nearby gene. It remains debated if such differences in mRNA levels correspond to changes in protein levels, since there is evidence of compensatory buffering at the translational level (Khan et al., 2013). It also remains unclear how differences in the level of a protein affects species fitness. In contrast, our results demonstrate that non-coding regions have the capacity to mediate an on/off switch of a conserved protein-coding gene.

KRAB-ZFPs are the largest family of transcription factors in the human genome and have rapidly expanded and evolved in the primate lineage. It has been proposed that KRAB-ZFPs are engaged in an arms race with TEs, in which KRAB-ZFPs evolve to bind specific TEs and silence their activity (Jacobs et al., 2014). In this model, TEs mutate their ZFP-targeted sites, thereby escaping from silencing and regaining activity. The KRAB-ZFP would then further evolve to once again silence the TE, resulting in a dynamic competition between KRAB-ZFPs and TEs that drive their evolution. This evolutionary mechanism has been proposed to have been used by host genomes to drive evolutionary mechanisms (Trono, 2016). In this study we provide evidence of the human-specific expression of several KRAB-ZFP transcription factors during brain development. One example, *ZNF558*, arose more than 100 million years ago to repress then-active LINE-1 transposons, but has since been co-opted to repress protein-coding genes. Although its TE targets have degenerated, ZNF558 is highly conserved among mammals, in line with functional biological roles independent of TE suppression. ZNF558 now binds to both ancient L1s – likely representing a genomic fossil – and several gene targets in human cells (Imbeault et al., 2017). During human brain development, we found that ZNF558 regulates a single protein-coding gene, *SPATA18*. The ZNF558-binding site in *SPATA18* is conserved only in primates, suggesting that ZNF558 may play other roles in other tissues to regulate more ancient co-opted targets. It would be interesting to chart the functional roles of ZNF558 in other more distantly-related mammals, where innovation in target sequences may have led to co-option of this transcription factor for other functional roles.

The genetic basis for the unusual on/off switch of ZNF558 expression between human and chimpanzee forebrain progenitors resides in a downstream variable number tandem repeat (VNTR). The length of the VNTR correlates with repression of the *ZNF558* locus: in humans, where the repeat is 30-40 units long, the locus is linked to active transcription while in non-human primates (NHPs) the VNTR tends to be much larger and this is associated with transcriptional repression. Our observations are in line with a model where the common ancestor of humans and NHPs carried a long VNTR downstream of *ZNF558*. This VNTR contracted after the human/chimp evolutionary split, likely due to positive selection, ultimately resulting in differences in SPATA18 levels and altered mitochondria homeostasis in the developing human brain.

We do not fully understand the molecular mechanisms that underlie how differences in VNTR unit copy number affects ZNF558 expression. We found that the VNTR is decorated by the repressive histone mark H3K9me3 both in human and chimpanzee fbNPCs. Since the chimpanzee VNTR is longer, a VNTR-based heterochromatin region is substantially larger. We also found evidence that the short human VNTR unit is transcribed, giving rise to a long non-coding RNA, which may influence in *cis* the expression of nearby genes such as *ZNF558*. The most likely explanation for the observed phenomenon is that the extended VNTR is a heterochromatic region and that its shortening reveals a brain-specific enhancer and/or lncRNA in the repeat. Such a copy-number dependent VNTR based epigenetic regulatory mechanism of nearby protein coding gene expression is reminiscent of what has previously observed at the *FSHD* locus (Cabianca et al., 2012). Our data indicate that the VNTR at the *ZNF558* locus is variable in the human population and that there are individuals with a repeat length similar to that of chimpanzee. Molecular and phenotypical analysis of such individuals, using e.g. iPSCs-based modeling, would be interesting and may be in line with the idea of broad roles for VNTRs in human phenotypical variation.

The downstream consequences of a shortened VNTR and activated ZNF558 expression is transcriptional repression of *SPATA18* in human fbNPCs. The product of *SPATA18* is MIEAP (mitochondrial eating protein), which clears mitochondria via mitophagy (Kitamura et al., 2011) as part of a homeostatic system. Emerging literature indicate that mitochondria are central in neural stem cell fate decisions and the process of neurogenesis partly through the control of a metabolic switch from glycolysis to OXPHOS occurring when NPCs differentiate to neurons (Arrázola et al., 2019; Khacho et al., 2016, 2019). ARHGAP11B, a human specific gene (Florio et al., 2015), is localized to the mitochondria where it induces a cancer-like metabolism to promote the proliferation of basal neural progenitors (Namba et al., 2020). This recent observation directly links mitochondrial function to speciation and the expansion of the human cerebral cortex (Namba et al., 2020). Exactly how *SPATA18*, regulated by ZNF558 uniquely in humans, feeds into this metabolic pathway, remains to be determined. However, it is notable that a range of human neurodevelopmental disorders including Williams Syndrome, Leigh Syndrome, autism spectrum disorders and schizophrenia have mitochondrial dysfunction as part of their reported phenotypes (Khacho et al., 2019; Tebbenkamp et al., 2018). We also note that DNA duplications of the *SPATA18* locus can lead to intellectual disability and delayed language and speech development (https://decipher.sanger.ac.uk), strengthening the notion that *SPATA18* and its partner transcription factor ZNF558 play important roles in human brain development.

In summary, our results illustrate how one KRAB-ZFP, ZNF558, became co-opted and subsequently contributed to human brain evolution. Future studies of KRAB-ZFPs and VNTRs in human evolution and human disease will be interesting and rewarding.

## Data availability

There are no restrictions in data availability. Accession code for the RNA and DNA sequencing data presented in this study is XXXXXX.

## Acknowledgements

We would like to thank Fred H. Gage, Wieland Huttner, Svante Pääbo, Barbara Treutlein, Steve Henikoff, He Zhisong, Mitchell Vollger and Yafei Mao for their helpful comments and computational and technical support. We also thank, M. Persson Vejgården and A. Hammarberg for technical assistance. We are grateful to all members of the Jakobsson lab. The work was supported by grants from the Swedish Research Council (2018-02694, JJ), the Swedish Brain Foundation (FO2019-0098, JJ), Cancerfonden (190326, JJ), Barncancerfonden (PR2017-0053, JJ), the Swedish Society for Medical Research (S19-01000, C.H.D.) and the Swedish Government Initiative for Strategic Research Areas (MultiPark & StemTherapy). E.E.E. is an investigator of the Howard Hughes Medical Institute.

## Author contributions

All authors took part in designing the study and interpreting the data. P.A.J, P.L.B. and J.J. conceived and designed the study. P.A.J., D.G., J.J., D.A.M.A., M.E.J. and K.P. performed experimental research. P.L.B, C.H.D., P.H, R. G., Y.S. and F.E. performed bioinformatic analyses. J.P., D.T. and E.E.E. contributed reagents and expertise. P.A.J, P.L.B, C.H.D. and J.J. wrote the manuscript and all authors reviewed the final version.

## MATERIALS AND METHODS

### iPSC culture

We used two human iPSC lines generated by mRNA transfection (RBRC-HPS0328 606A1 and RBRC-HPS0360 648A1, both from RIKEN; from here on referred to as HS1 and HS2, respectively). We used two chimpanzee iPSC lines: one generated by mRNA transfection (Sandra A, herein referred to as PT1) and the other with viral vector transduction (PR00818 PTCL-5, herein referred to as PT2) (Marchetto et al., 2013; Mora-Bermúdez et al., 2016). iPSCs were maintained on LN521-coated (0.7 µg/cm^2^; Biolamina) Nunc multidishes in iPS media (StemMACS iPS-Brew XF and 0.5% penicillin/streptomycin (Gibco)). Cells were passaged 1:2-1:6 every 2-4 days by being rinsed once with DPBS (Gibco) and dissociated using 0.5 mM EDTA (75 µl/cm^2^; Gibco) at 37 °C for 7 minutes. Following incubation, EDTA was carefully aspirated from the well and the cells were washed off from the dish using washing medium (9.5 ml DMEM/F-12 (31330-038; Gibco) and 0.5 ml knockout serum replacement (Gibco)). The cells were then centrifuged at 400*g* for 5 minutes and resuspended in iPS brew medium supplemented with 10 µM Y27632 (Rock inhibitor; Miltenyi) for expansion. The media was changed daily (Grassi et al., 2020; Nolbrant et al., 2017).

### Differentiation into forebrain neural progenitors (fbNPCs)

iPSCs were grown to approximately 70-90% confluency and were then dissociated as usual for passaging. After centrifugation, the cells were resuspended in N2 medium (1:1 DMEM/F-12 (21331-020; Gibco) and Neurobasal (21103-049; Gibco) supplemented with 1% N2 (Gibco), 2 mM L-glutamine (Gibco), and 0.2% penicillin/streptomycin). The cells were manually counted twice and plated at a density of 10,000 cells/cm^2^ in 250 µl medium/cm^2^ on LN111 Nunc Δ multidishes (1.14 µg/cm^2^; Biolamina). 10 µM SB431542 (Axon) and 100 ng/ml noggin (Miltenyi) for dual SMAD inhibition, and 10 µM Y27632 was added to the medium. The medium was changed every 2-3 days (N2 medium with SB431542 and noggin) until day 9 of differentiation, when N2 medium without SMAD inhibitors was used. On day 11, the cells were replated by washing twice with DPBS followed by adding StemPro accutase (75 µl/cm^2^; Gibco) for 10-20 minutes at 37 °C. The dissociated cells were washed off with 10 ml wash medium, centrifuged for 5 minutes at 400*g* and resuspended in B27 medium (Neurobasal supplemented with 1% B27 without vitamin A (Gibco), 2 mM L-glutamine and 0.2% penicillin/streptomycin Y27632 (10 µM), BDNF (20 ng/ml; R&D), and L-ascorbic acid (0.2 mM; Sigma). The cells were counted and replated at 800,000 cells/cm^2^ on LN111-coated plastic in B27 medium (600 µl medium/cm^2^). The cells were kept in the same medium until day 14, after which new B27 medium was added (Grassi et al., 2020).

### Immunocytochemistry

The cells were washed once with DPBS and fixed for 15 minutes with 4% paraformaldehyde (Merck Millipore), followed by three rinses with DPBS. The fixed cells were then pre-blocked for a minimum of 30 minutes in a blocking solution of KPBS with 0.25% triton-X100 (Fisher Scientific) and 5% donkey serum. The primary antibody (rabbit anti-FOXG1, 1:50 dilution, Abcam, RRID: AB_732415 and anti-NANOG, 1:100 dilution, Abcam, RRID: AB_446437) was added and incubated overnight. On the following day, the cells were washed twice with KPBS. The secondary antibody (donkey anti-rabbit Cy3; 1:200; Jackson Lab) was added with DAPI (1:1000; Sigma-Aldrich) as a nuclear counterstain and incubated at room temperature for one hour, followed by 2-3 rinses with KPBS. The cells were visualized on a Leica microscope (model DMI6000 B), and images were cropped and adjusted in Adobe Photoshop CC.

### Bulk RNA sequencing

On the day of harvest, the cells were washed once with PBS and lysed with 350 µl RLT buffer with 1% mercaptoethanol (Thermo Fisher). RNA was extracted using the RNeasy mini kit (Qiagen) according to manufacturer’s protocol. The quality and concentration of the RNA was analyzed using 2100 Bioanalyzer (RNA nano; Agilent) and Qubit (RNA HS assay kit). Libraries for sequencing were prepared using the TruSeq RNA Library Prep kit v2 (Illumina) and again quality-controlled using the Bioanalyzer (high-sensitivity DNA assay) and Qubit (dsDNA HS assay kit). Finally, the libraries were sequenced using an Illumina NextSeq 500, 150-bp paired-end reads (300 cycles).

RNA-sequencing samples were mapped to the human reference genome (GRCh38) and the chimp reference genome (Clint_PTRvs2/PanTro6) using STAR aligner v2.5.0 (Dobin et al., 2013), allowing 0.03 mismatches per base (--outFilterMismatchNoverLmax 0.03), using Gencode v27 (Harrow et al., 2012) gene models for splice junction annotation. For multimapping, STAR was run on default settings, retaining reads that map at up to 10 loci. For unique mapping, STAR was run with --outFilterMultimapNmax 1. Gene counts were quantified with the Subread package FeatureCounts (Liao et al., 2014), counting reads overlapping Gencode (v27) gene annotations. For mapping and quantification of chimp samples, the Gencode annotation for GRCh38 were lifted to the panTro6 reference genome using the UCSC LiftOver tool. Normalization and differential expression analysis was performed with the R package DESeq2 (Love et al., 2014).

To analyse TE expression, we mapped the samples once again using STAR aligner v2.6.0 (Dobin et al., 2013), allowing reads to map 100 times (--outFilterMultimapNmax 100) with 200 anchors (--winAnchorMultimapNmax 200). The bam files were input to TEtranscripts version 2.0.3 (Jin et al., 2015) in multi-mode (--mode multi) using gencode annotation GRCh38 as the gene annotation (--GTF), as well as the curated GTF file of TEs provided by TEtranscripts’ authors (--TE). Differential expression analysis was performed using DESeq2 (Love et al., 2014).

### Single-cell RNA sequencing

HS1 and PT1 were differentiated to day 14, washed twice with DPBS, and dissociated with Accutase for approximately 10 minutes, followed by centrifugation at 400 × g for 5 minutes in wash medium. All pipetting was done very gently to avoid cell death. The pellets were resuspended in 1 ml PBS with 0.04% BSA (Sigma) and filtered through 100 µm cell strainers (Falcon) twice. Cells were resuspended in order to yield a concentration of approximately 1000 cells/µl. The single-cell libraries were prepared with Chromium Single Cell A chip kit and Chromium Single Cell 3’ Library & Gel Bead kit v2 (10× Genomics), quality controlled, and quantified using Qubit ds DNA HS and Bioanalyzer High Sensitivity DNA Assay prior to sequencing. The samples were sequenced for 26 cycles on read 1 and 98 cycles on read 2 using the Illumina NextSeq 500. Raw single-cell RNA-seq data was processed using the Cell Ranger software suite. Raw base call files were converted using cellranger mkfastq before aligning, filtering, barcode count, and UMI counting was performed using cellranger count. Count matrices were further analyzed using the Seurat R package (Butler et al., 2018). The data were filtered on number of genes detected in each cell (2000-6000 genes/cell in human and 2000-5000 genes/cell in chimpanzee were kept for further analysis), and only cells with max 0.05% mitochondrial gene reads. 4553 HS1 cells were sequenced and 4355 were used in the analysis, 5674 PT1 cells were sequenced and 3620 were kept after filtering. The data was further log2-normalized and scaled to total expression in each cell, before further scaling cells on number of UMIs detected and percentage of mitochondrial gene count. PCA was run on variable genes defined using the FindVariableGenes function. tSNE was run using PCA dimensionality reduction.

### Lentiviral production

Lentiviral vectors were produced according to Zufferey et al. (Zufferey et al., 1997) and were in titers of 10^8^ – 10^9^ TU/ml as determined by qRT-PCR. Briefly HEK293T cells were grown to a confluency of 70 – 90 % at the day of transfection for lentiviral production. Third-generation packaging and envelop vectors (pMDL, psRev, and pMD2G) together with Polyethyleneimine (PEI Polysciences PN 23966, in DPBS (Gibco) were used. The lentivirus was harvested two days after transfection The supernatant was then collected, filtered and centrifuged at 25,000*g* for 1.5 hours at 4 °C. The supernatant was removed from the tubes and the virus was resuspended in PBS and left at 4 °C. The resulting lentivirus was aliquoted and stored at −80 °C.

### FACS

Cells were detached with Accutase, resuspended in differentiation media with Rock inhibitor (10µM, Miltenyi) and Draq7 (BD Bioscience), and strained (70µm, BD Bioscience). Gating parameters were determined by side and forward scatter to eliminate debris and aggregated cells. The GFP-positive gates were set using untransduced cells. The sorting gates and strategies were validated via reanalysis of sorted cells (>95% purity cut-off). GFP-positive/Draq7-negative single cells (100,000-200,000 cells/pellet) were collected, spun down (400*g*, 10 min) and snap frozen on dry ice. Cell pellets were kept at −80°C until RNA or DNA was isolated.

### CUT & RUN

We followed the protocol detailed by the Henikoff lab (Skene et al., 2018). Briefly, 100,000 cells were washed twice (20 mM HEPES pH 7.5, 150 mM NaCl, 0.5 mM spermidine, 1x Roche cOmplete protease inhibitors) and attached to 10 ConA-coated magnetic beads (Bangs Laboratories) that had been pre-activated in binding buffer (20 mM HEPES pH 7.9, 10 mM KCl, 1 mM CaCl_2_, 1 mM MnCl_2_). Bead-bound cells were resuspended in 50 µL buffer (20 mM HEPES pH 7.5, 0.15 M NaCl, 0.5 mM Spermidine, 1x Roche complete protease inhibitors, 0.02% w/v digitonin, 2 mM EDTA) containing primary antibody (rabbit anti-H3K9me3, Abcam ab8898; rabbit anti-H3K27me3, Cell Signaling Technology C36B11; rabbit anti-H3K4me3 Active Motif 39159; or goat anti-rabbit IgG, Abcam ab97047) at 1:50 dilution and incubated at 4 °C overnight with gentle shaking. Beads were washed thoroughly with digitonin buffer (20 mM HEPES pH 7.5, 150 mM NaCl, 0.5 mM Spermidine, 1x Roche cOmplete protease inhibitors, 0.02% digitonin). After the final wash, pA-MNase (a generous gift from Steve Henikoff) was added in digitonin buffer and incubated with the cells at 4 °C for 1 h. Bead-bound cells were washed twice, resuspended in 100 µL digitonin buffer, and chilled to 0-2 °C. Genome cleavage was stimulated by addition of 2 mM CaCl2 at 0 °C for 30 min. The reaction was quenched by addition of 100 µL 2x stop buffer (0.35 M NaCl, 20 mM EDTA, 4 mM EGTA, 0.02% digitonin, 50 ng/µL glycogen, 50 ng/µL RNase A, 10 fg/µL yeast spike-in DNA (a generous gift from Steve Henikoff)) and vortexing. After 10 min incubation at 37 °C to release genomic fragments, cells and beads were pelleted by centrifugation (16,000 g, 5 min, 4 °C) and fragments from the supernatant purified. Illumina sequencing libraries were prepared using the Hyperprep kit (KAPA) with unique dual-indexed adapters (KAPA), pooled and sequenced on a Nextseq500 instrument (Illumina). Paired-end reads (2×75) were aligned to the human and yeast genomes (hg38 and R64-1-1 respectively) using bowtie2 (--local –very-sensitive-local –no-mixed –no-discordant –phred33 -I 10 -X 700) and converted to bam files with samtools (Langmead and Salzberg, 2012; Li et al., 2009). Normalized bigwig coverage tracks were made with bamCoverage (deepTools) (Ramírez et al., 2014), with a scaling factor accounting for the number of reads arising from the spike-in yeast DNA (10^4/aligned yeast read number). Tracks were displayed in IGV.

### ChIP-exo analysis

ZNF558 and ZNF557 ChIP-exo data was downloaded along with input control from <u>GSE78099</u> (Imbeault et al., 2017). Raw ChIP-exo reads were quality controlled with FastQC (Babraham) and aligned to the human reference genome (GRCh38) using bowtie2 with -- sensitive-local (Langmead and Salzberg, 2012). Unique reads were filtered by retaining only alignments with MAPQ>10 (samtools view -q 10) (Li et al., 2009). Peaks were called with HOMER (Heinz et al., 2010) and those with score >20 retained. Visualization of ChIP signals was done in deepTools using the computeMatrix and plotHeatmap modules (Ramírez et al., 2014). ZNF558 motif analysis was done with MEME (search for one motif, length 18-30 nt) (Bailey et al., 2015). Prediction of the ZNF558 DNA binding sequence was done as detailed in Persikov & Singh (Persikov and Singh, 2014).

### CRISPRi

In order to silence the transcription of ZNF558 we used the catalytically inactive Cas9 (deadCas9) fused to the transcriptional repressor KRAB. Single guide sequences were designed to recognize DNA regions just down-stream of the transcription start site (TSS) according to the GPP Portal (Broad Institute). See Table 1 for guide RNA sequences. For the induction of repressive marks on the human VNTR we chose three guide RNAs from the UCSC Gene Browser CRISPR target track with high on-target specificity and only found within the VNTR region and avoiding transposable elements (RepeatMasker). See Table 1 for guide RNA sequences. The guides were inserted into a deadCas9-KRAB-T2A-GFP lentiviral backbone containing both the guide RNA under the U6 promoter and dead-Cas9-KRAB and GFP under the Ubiquitin C promoter (pLV hU6-sgRNA hUbC-dCas9-KRAB-T2a-GFP, a gift from Charles Gersbach, Addgene plasmid #71237 RRID:Addgene_71237). The guides were inserted into the backbone using annealed oligos and the BsmBI cloning site. Lentiviruses were produced as described below yielding titers between 4.9E+08 and 9.3E+09. Control virus with a gRNA sequence not present in the human genome (LacZ) was also produced and used in all experiments. All lentiviral vectors were used with an MOI between 5 and 20. Cells were FACS sorted as described above and knock-down efficiency was validated using standard quantitative real-time RT-PCR techniques.

**Table 1.**
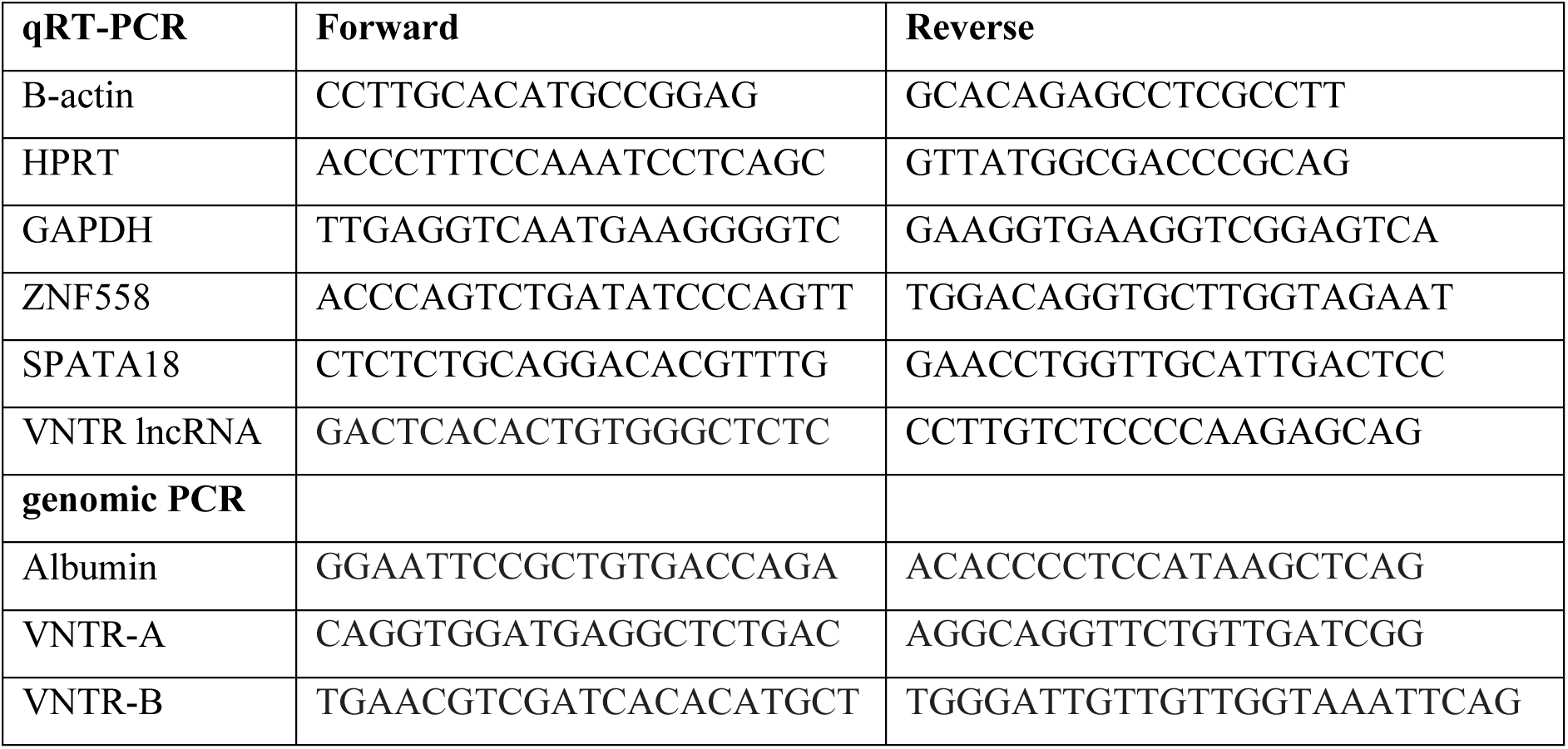
Primer sequence for PCR

ZNF558-g1 GCCAAAAGCGCCGACTCGCG; ZNF558-g2 AGTCGGCGCTTTTGGCCCCG; LacZ; TGCGAATACGCCCACGCGAT; VNTR-g1 GCTGCCCTGAGATATGTGTG; VNTR-g2 TACTGGAATGGGTAGGAATG; VNTR-g3 CCAGGAGCTGCACATTGAGG

### ZNF558 overexpression in chimpanzee cells

Chimpanzee cells (PT1) was transduced with a lentiviral construct overexpressing ZNF558 and the puromycin resistance cassette. An identical virus without the ZNF588 insert was used as a control. Cells were split and transduced and allowed to recover for 2-3 days,followed by puromycin selection. The cells were in puromycin selection for more than 5 days, then expanded briefly without puromycin and frozen. Cells were thawed and exposed to an additional 1-2 day of puromycin selection prior to start of differentiation. Puromycin was added to the differentiation media at day 0, day 4, day 7 and day 11.

### qRT-PCR

Total RNA was first extracted according to the supplier’s recommendations using the mini RNeasy kit (Qiagen). cDNA was generated using the Maxima First Strand cDNA Synthesis Kit (Thermo Scientific) and analysed with SYBR Green I master (Roche) on a LightCycler 480 (Roche). Data are represented with the ΔΔCt method normalized to the housekeeping genes B-actin, GAPDH and HPRT1. See Primer sequences are listed in Table 1. Expression levels were confirmed using one additional primer pair (data not shown).

### Copy number analysis and mitochondrial analysis

Genomic and mitochondrial DNA was extracted using DNeasy Blood and Tissue Kits (Qiagen). All primers were used together with LightCycler 480 SYBR Green I Master (Roche). For mitochondrial analysis we used previously published primers used for mitochondrial copy number assessment (MtDNA; FP CACCCAAGAACAGGGTTTGT, RP TGGCCATGGGTATGTTGTTA, NucDNA FP TGCTGTCTCCATGTTTGATGTATCT, RP TCTCTGCTCCCCACCTCTAAGT (Rooney et al., 2015).

For VNTR unit copy number assessment we designed primers for different parts of the VNTR repeat unit that was not found in other parts of the genome. We used albumin as a positive control with the albumin copy number set as 2 (see Table 1)

### Evolutionary analysis of ZNF558

Phylogenetic tree, pairwise sequence alignment scores and dN/dS ratios of ZNF558 orthologues was downloaded from the Ensembl (Zerbino et al., 2018). Orthologues section for ZNF558 in nexus format and FigTree was used for visualization (http://tree.bio.ed.ac.uk/software/figtree/). For analysis of DNA binding residues of the zinc finger domain of ZNF558 and orthologues, the ClustalW multiple sequence alignment from Ensembl orthologues was visualized in JalView. DNA contacting residues were defined the four amino acids in position −1, −4, −5 and −7 relative to the first histidine residue of the two zinc-coordinating histidine residues. For analysis of ZNF558 TE binding, TE coordinates were downloaded from RepeatMasker.

### Copy number estimation for the VNTR locus near ZNF558 in human and non-human great ape samples

To investigate the copy number variation for the large VNTR downstream of *ZNF558* in human and non-human great ape lineages, we applied a read-depth based copy number genotyper (Sudmant et al., 2010) to a collection of 1,112 high-coverage genomes from several publicly-available resources (Bergström et al., 2020; Mafessoni et al., 2020; Mallick et al., 2016; Meyer et al., 2012; Prado-Martinez et al., 2013; Prüfer et al., 2017). In short, sequencing reads were divided into multiples of 36-mer, which were then mapped to a repeat-masked human reference genome (GRCh38) using mrsFAST (Hach et al., 2010). Up to two mismatches per 36-mer were allowed in order to increase our mapping sensitivity and read depth of mappable sequences in our analysis was corrected for underlying GC content. Finally, copy number estimate for the locus of interest was computed by summarizing over all mappable bases for each sample.

## Supplementary Figures

**Fig S1.**
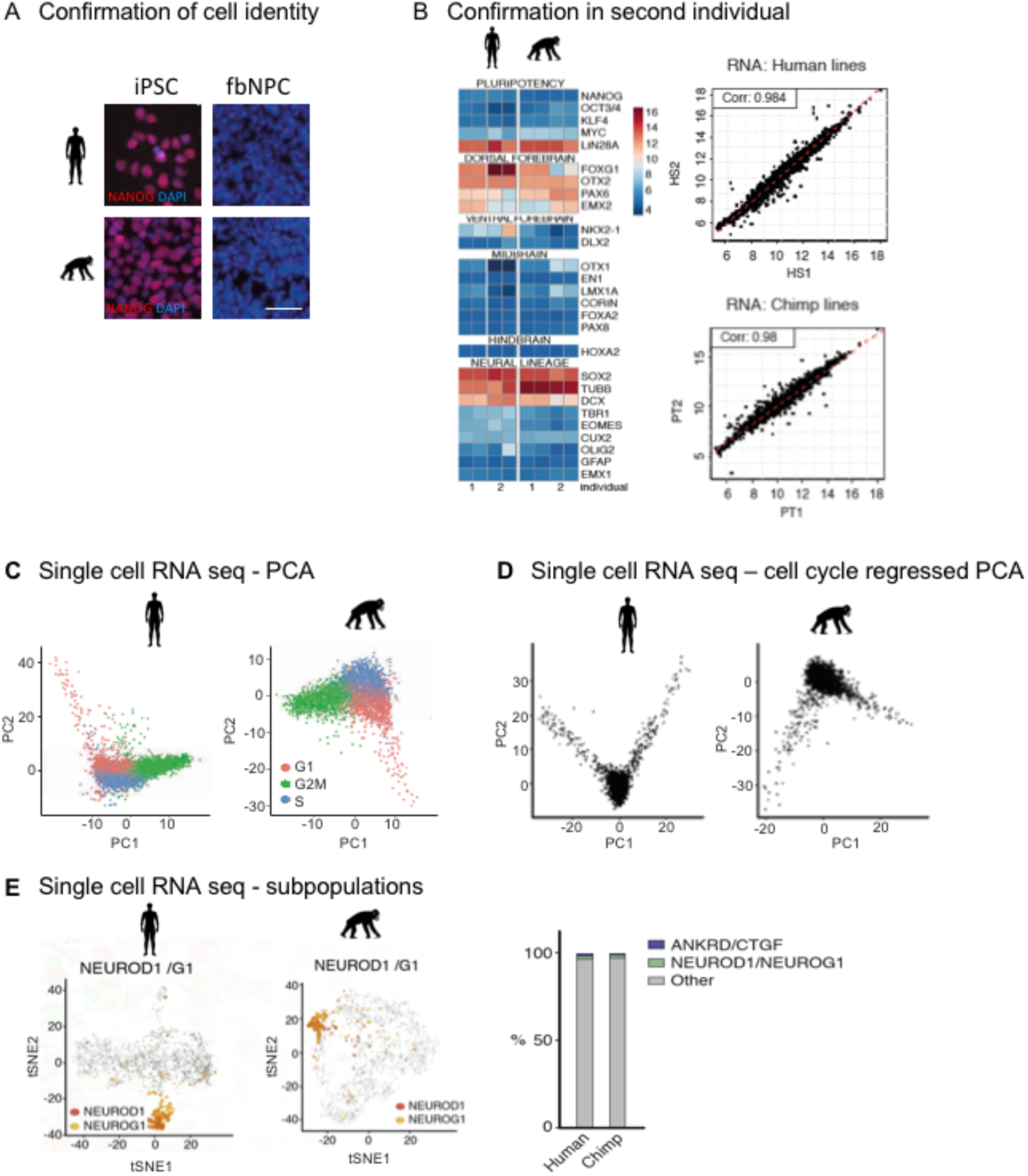
Confirmation of cell identity. (A) Nanog immunocytochemistry in human and chimp iPSCs and differentiated fbNPCs. Nuclei stained with DAPI (blue). The scale bar represents 50mm. (B) Heatmap of neural marker transcript expression at day 14 of differentiation in two individuals of each species. (C) Principal component analysis of single-cell RNA-seq divided into groups based on cell-cycle stages. (D) Principal component analysis of single-cell RNA-seq, where cell-cycle effects have been regressed out. (E) Left: tSNE of human and chimpanzee single-cell RNA-seq data showing the expression of the neuronal markers NEOROD1/G1. Right: Bar charts showing the low percentage of ANKRD1/CTGF+ and NEUROD1/NEUROG1+ cells in single-cell RNA-seq.

**Fig S2.**
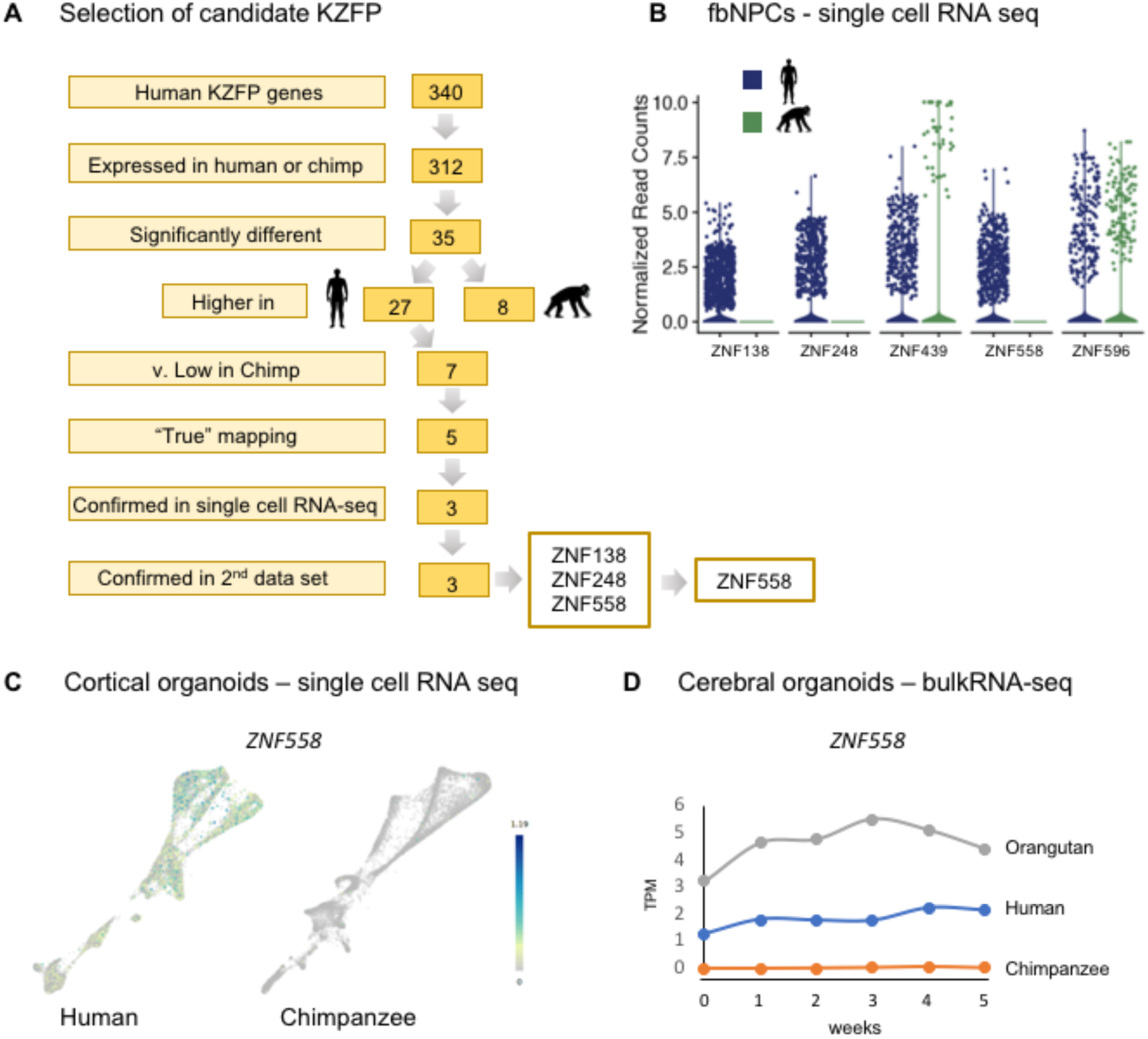
Selection and validation of KZFP candidates. (A) Flow-diagram outlining the selection process in finding KZFP candidates expressed only in human fbNPCs. (B) Normalized read counts of each KZFP candidate in single cell sequencing data of human (blue) and chimpanzee (green) fbNPCs. Each dot represents one cell. (C) ZNF558 expression in single cell RNA-seq analysis from cortical organoids. Data from Kanton et al., 2019. (D) ZNF558 expression in bulkRNA-seq from cerebral organoids. TPM = transcript per million. Data from Field et al 2019.

**Fig S3.**
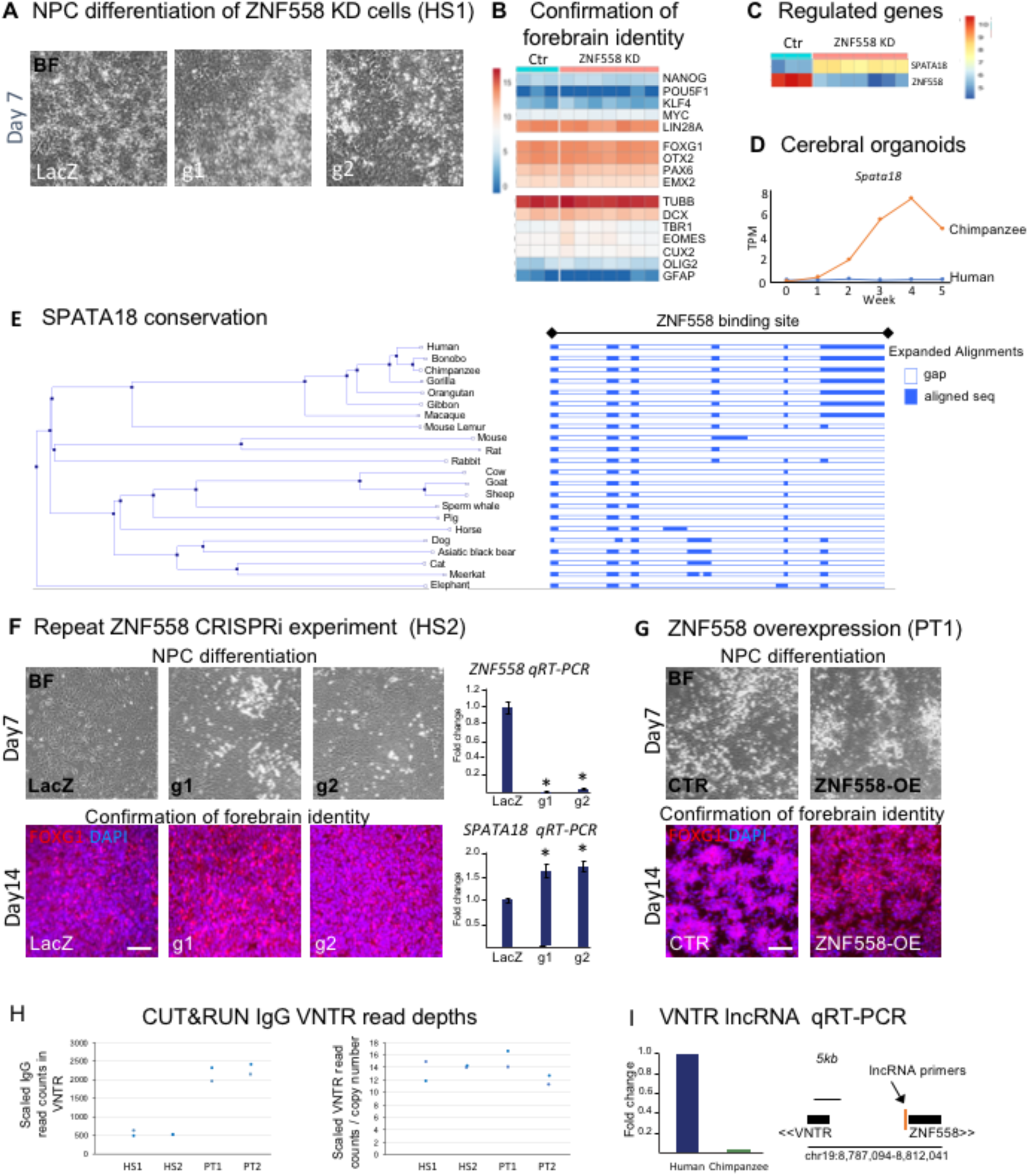
ZNF558 binds and regulates SPATA18. (A-C) CRISPRi mediated repression of ZNF558. Differentiation (A) and confirmation of fbNPC forebrain identity at Day14 (B). (C) Heatmap of all differentially-regulated genes after ZNF558 inhibition. (D) SPATA18 expression in bulkRNA-seq from cerebral organoids. TPM, transcripts per million. Data from Field et al 2019. (E) Multiple sequence alignment of the ZNF558 binding site in *SPATA18*, showing conservation in primates. Graphics were retrieved from Ensembl comparative genomics alignment webtool. (F) CRISPRi mediated repression of ZNF558 in HS2, confirming SPATA18 up-regulation (bottom-right. n=4, * = p<0.05 (Student’s t-test), data is represented as mean +/− s.e.m. Scale bar = 50mm, (G) Differentiation of ZNF558-overexpressing chimpanzee cells (top) and the confirmation of forebrain identity at Day14 using FOXG1 immunocytochemistry (bottom). Scale bar = 50mm. (H) CUT&RUN IgG read depths arising from VNTR (hg38 chr19: 8,737,106-8,790,930) without (left) and with (right) additional scaling factor for individual VNTR copy number. Shown are two technical replicates from two biological replicates (different individuals) (I) qRT-PCR of a lncRNA transcript originating in the VNTR, n=2 (left) and location of the primer pairs (left). BF, brightfield microscopy.

